# Metabolic clearance rate modeling: A translational approach to quantifying cerebral metabolism using hyperpolarized [1-13C]pyruvate

**DOI:** 10.1101/2022.11.02.514924

**Authors:** James T. Grist, Nikolaj Bøgh, Esben Søvsø Hansen, Anna M. Schneider, Richard Healicon, Vicky Ball, Jack J.J.J. Miller, Sean Smart, Yvonne Couch, Alastair Buchan, Damian J. Tyler, Christoffer Laustsen

## Abstract

Hyperpolarized carbon-13 MRI is a promising technique for *in vivo* metabolic interrogation of alterations between health and disease. This study introduces a model-free formalism for quantifying the metabolic information in hyperpolarized imaging.

This study investigated a novel model-free perfusion and metabolic clearance rate (MCR) model in pre-clinical stroke and in the healthy human brain.

Simulations showed that the proposed model was robust to perturbations in T_1_, transmit B_1_, and k_PL_. A significant difference in ipsilateral vs contralateral pyruvate derived cerebral blood flow (CBF) was detected in rats (140 ± 2 vs 89 ± 6 mL/100g/min, p < 0.01, respectively) and pigs (139 ± 12 vs 95 ± 5 mL/100g/min, p = 0.04, respectively), along with an increase in fractional metabolism (26 ± 5 vs 4 ± 2 %, p < 0.01, respectively) in the rodent brain. In addition, a significant increase in ipsilateral vs contralateral MCR (0.034 ± 0.007 vs 0.017 ± 0.02 s^-1^, p = 0.03, respectively) and a decrease in mean transit time (MTT) (31 ± 8 vs 60 ± 2, p = 0.04, respectively) was observed in the porcine brain. In conclusion, MCR mapping is a simple and robust approach to the post-processing of hyperpolarized magnetic resonance imaging.

## Introduction

The clinical translation of dissolution dynamic nuclear polarization hyperpolarized magnetic resonance (MR) is currently ongoing at sites around the world^1–6^. The method has already shown its value in the pre-clinical setting, while the clinical value of this technology is in the process of being demonstrated^7–14^. Previous studies have proposed apparent rate constants^15,16^, time-to-peak^17^, and area-under-the-curve^18^ as quantitative surrogates for apparent metabolic activity. One of the most popular approaches to quantifying data is to divide the summed signal from a metabolite (for example [1-13C]lactate, by the total observed substrate, for example, [1-13C]pyruvate). In the study, a model-free approach, currently used in both ^1^H perfusion MRI and in Positron Emission Tomography (PET), is applied to hyperpolarized imaging to provide a complementary quantifiable measure of the hyperpolarized data. Indeed, one major obstacle to clinical translation is the lack of general consensus on the acquisition and analysis of hyperpolarized data. This is due, in part, to the nature of the signal being a complicated mixture of unknown origin, and as such, it is difficult to develop quantifiable measures that allow general comparison across sites and species^19^. The approach used in this study is based upon commercially available software, allowing for the rapid introduction of the method into clinical studies, and may help tackle this challenge.

The general premise behind the approach used in this study is that the passage of a tracer injected into the body can be tracked through time, and the subsequent change in MRI signal post-processed to estimate quantitative perfusion-related parameters. These parameters include: the volume of tracer in a given location (sometimes referred to as ‘Cerebral Blood Volume’ or CBV), flow speed of the tracer (Cerebral Blood Flow or CBF), time taken to reach peak concentration (TTP), the mean time that the tracer spends in the vascular system (MTT), the signal decay (T_1_) and the rate of metabolic consumption of the tracer^20^. In the case of a non-metabolized, non-hyperpolarized tracer, such as gadolinium, the last two terms are ignored.

### The metabolic clearance rate

The metabolic clearance rate (MCR) can be defined as the amount of tracer that leaves a given tissue, either by metabolic conversion or by flow. The MCR parameter is used in positron emission tomography (PET) imaging, where the effective decay of radioactive signal is attributed to the metabolic conversion of the tracer of interest^21^. MCR has been shown to correlate with myocardial and renal oxidative metabolism ^22,23^ and, as such, offers a quantitative measure of the metabolic status of tissue *in vivo*.

An approach for determining the metabolic clearance rate has been demonstrated, which utilizes the mean transit time (MTT) of the metabolic tracer concentration (C(t)) in tissue^24^:

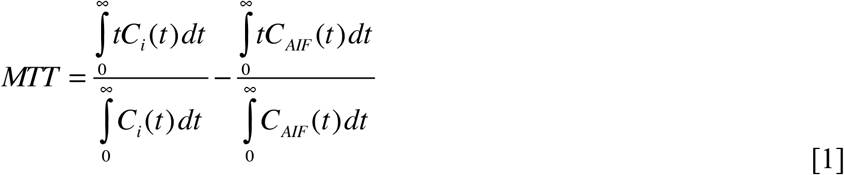

Where C_*i*_ is the signal in a voxel at a given time, and C_*AIF*_ is the signal of an arterial input function free assumed from metabolism. Thus, the fraction of [1-13C]pyruvate turnover in a hyperpolarized experiment can be written as a linear combination of the contribution of perfusion and the metabolic decay.

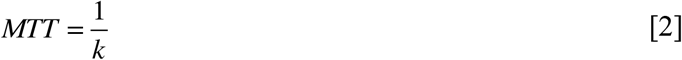

Where *k* is the metabolic clearance rate (s^-1^). It is known that estimations of the MTT of a hyperpolarized tracer need to be corrected for radiofrequency (RF) decay^24^. However, the calculation of the flow of a hyperpolarized tracer through a voxel is independent of RF decay. As a hyperpolarized experiment relies on supraphysiological concentrations of tracer, and under the assumption that the intracellular levels of pyruvate are small, it can be assumed that intracellular pyruvate is very small in comparison to the large extracellular signal. Therefore, [1-13C]pyruvate may be considered a “surrogate” first-pass perfusion marker with limited metabolic conversion of the pyruvate pool in the blood. Thus, it can be assumed that the MTT of pyruvate is approximately that of a gadolinium or arterial spin labelling experiment. Thus, if the estimated MTT for [1-13C]pyruvate and from a perfusion experiment are not equal, this implies that metabolism is affecting the apparent passage of the tracer. Therefore, a final extension to [2] is to combine the *k* _Gadolinium_ and *k*_Pyruvate_ to produce ‘Metabolic Clearance Rate’ (MCR) maps:

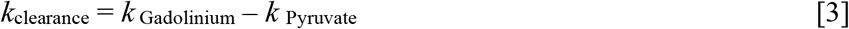

However, as the MTT in a hyperpolarized experiment needs to be corrected for T_1_-mediated signal decay^24^, which can be confounded with the many complexities of understanding transmit B_1_ inhomogeneity and regions variations in T_1_^25^, it may be easier to consider the blood flow differences between a gadolinium and pyruvate experiment, given that flow (mL/100g/min) is not confounded by RF inhomogeneity.

Considering the contribution of perfusion and metabolism is particularly important in situations with reduced blood flow, there may be cases where metabolism is maintained or changed^26^. If CBF_Pyruvate_ ≠CBF_Gadolinium_, this implies that metabolism is affecting the pyruvate signal.

Therefore, combining CBF_Gadolinium_ and CBF_Pyruvate_ could be used to determine the relative contribution of metabolism and perfusion to the pyruvate signal via:

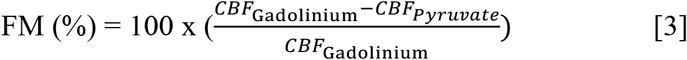

Where FM is termed the ‘fractional metabolism’, CBF_Gadolinium_ and CBF_Pyruvate_ are the Cerebral Blood Flow measurements of the gadolinium derived and pyruvate perfusion imaging, respectively.

This study combined data from pre-clinical small and large animal experiments, and human hyperpolarized imaging to demonstrate the potential use of metabolic clearance rate mapping in the brain.

## Results

### Simulation analysis

An overview of the methodology used in this study is shown in Figure 1, with an example of simulated curves for pyruvate and lactate shown in Supporting Figure S1. Simulation perturbations revealed differing effects upon the final estimated CBV/CBF/MTT depending on the signal-to-noise ratio (SNR), radiofrequency transmit inhomogeneity (B_1_^+^), [1-13C]pyruvate T_1_, and metabolic consumption rate (*k*_PL_) assumed.

**Figure 1.**
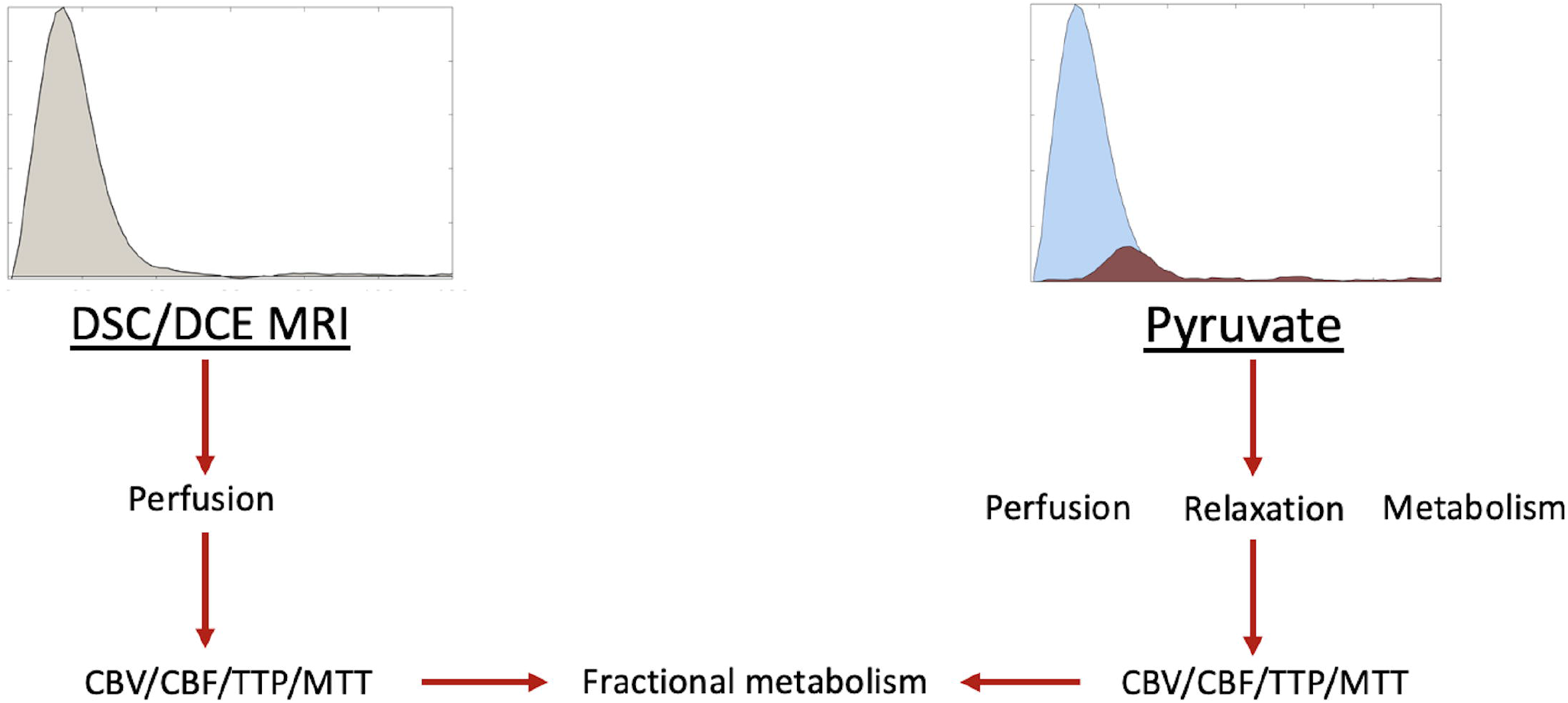
an example overview of the processing used in this methodology.

Perturbing [1-13C]pyruvate SNR showed minimal alterations in the calculated CBV, CBF, and MTT – with results showing less than 10% error at an SNR as low as 6 in all parameters (see Figure 2). Further perturbations to the transmit B_1_ error at a low flip angle (10 degrees) show that this method is very stable for CBV. However, CBF, MTT, and MCR were affected by B1+ deviation from the nominal flip angle (see Figure 3), with the maximum deviation being 8%.

**Figure 2.**
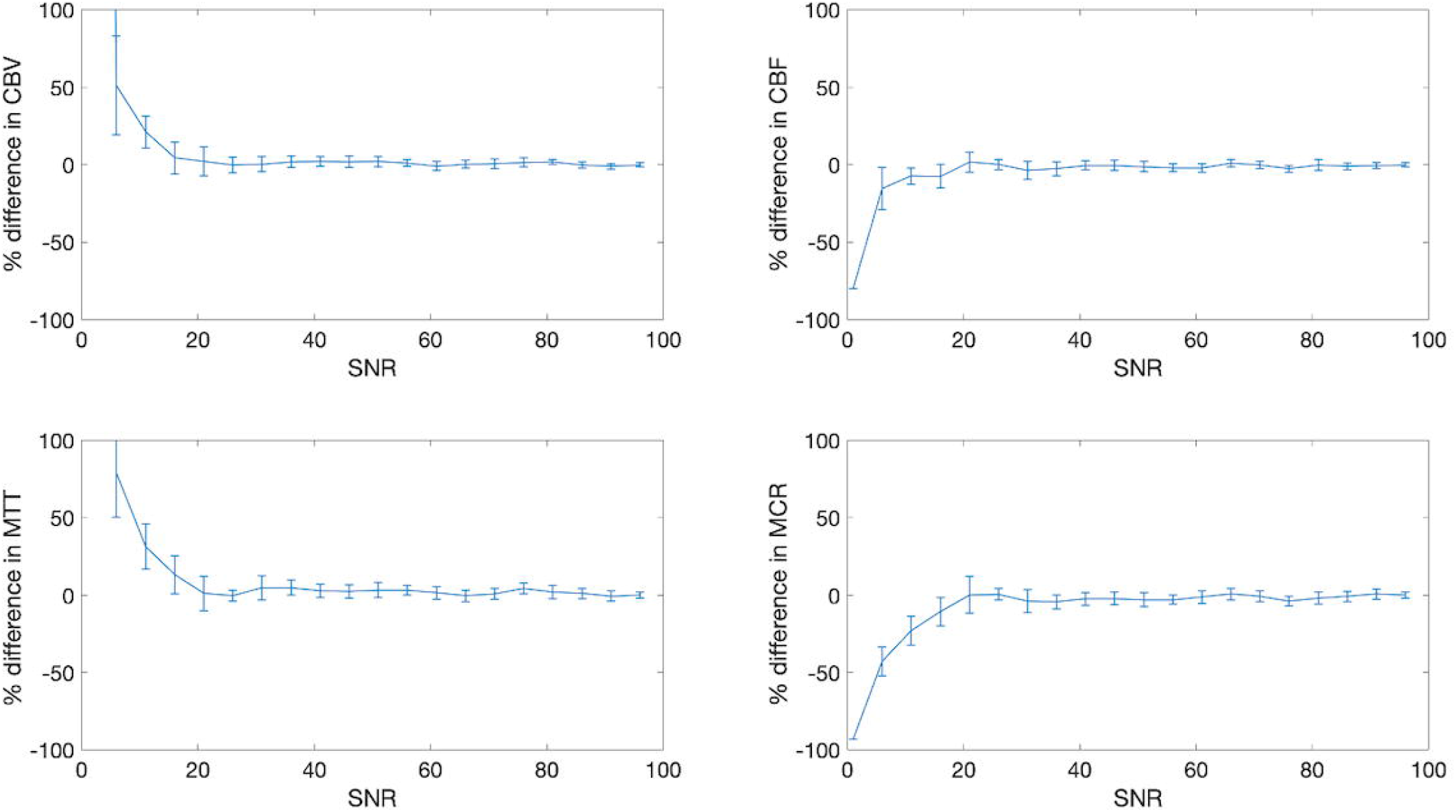
Simulation results demonstrating relative model stability down to SNR of 15 for all parameters. T_1_pyruvate, k_PL_, flip angle, and B_1_ error are assumed to be 35s, 0, 10 degrees, and 1, respectively.

**Figure 3.**
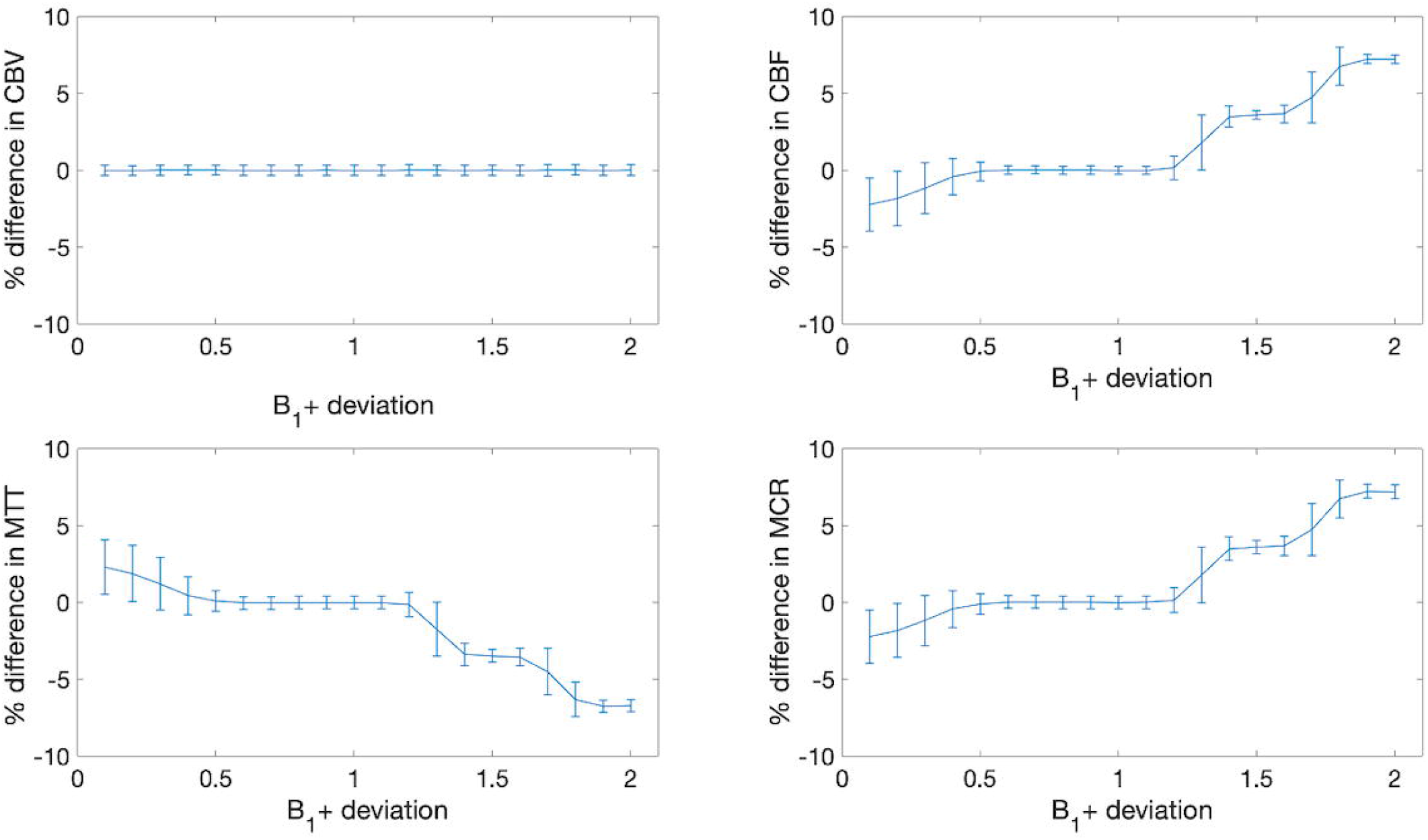
Simulation results demonstrating relative model stability for all parameters due to B_1+_ deviation. T_1_pyruvate, k_PL_, flip angle, and SNR are assumed to be 35s, 0, 10 degrees, and 100, respectively.

Varying k_PL_ revealed a subsequent decrease in CBV, CBF, and MTT – with a maximum difference of −28%, −10%, and −20% for CBV, CBF and MTT, respectively. MCR rose with increasing k_PL_, to a maximum increase of 27% at k_PL_ = 0.02 s^-1^, graphically shown in Figure 4. Varying the assumed T_1_ of [1-13C]pyruvate revealed that the CBF, MTT, and MCR varied approximately linearly, as shown in Figure 5.

**Figure 4.**
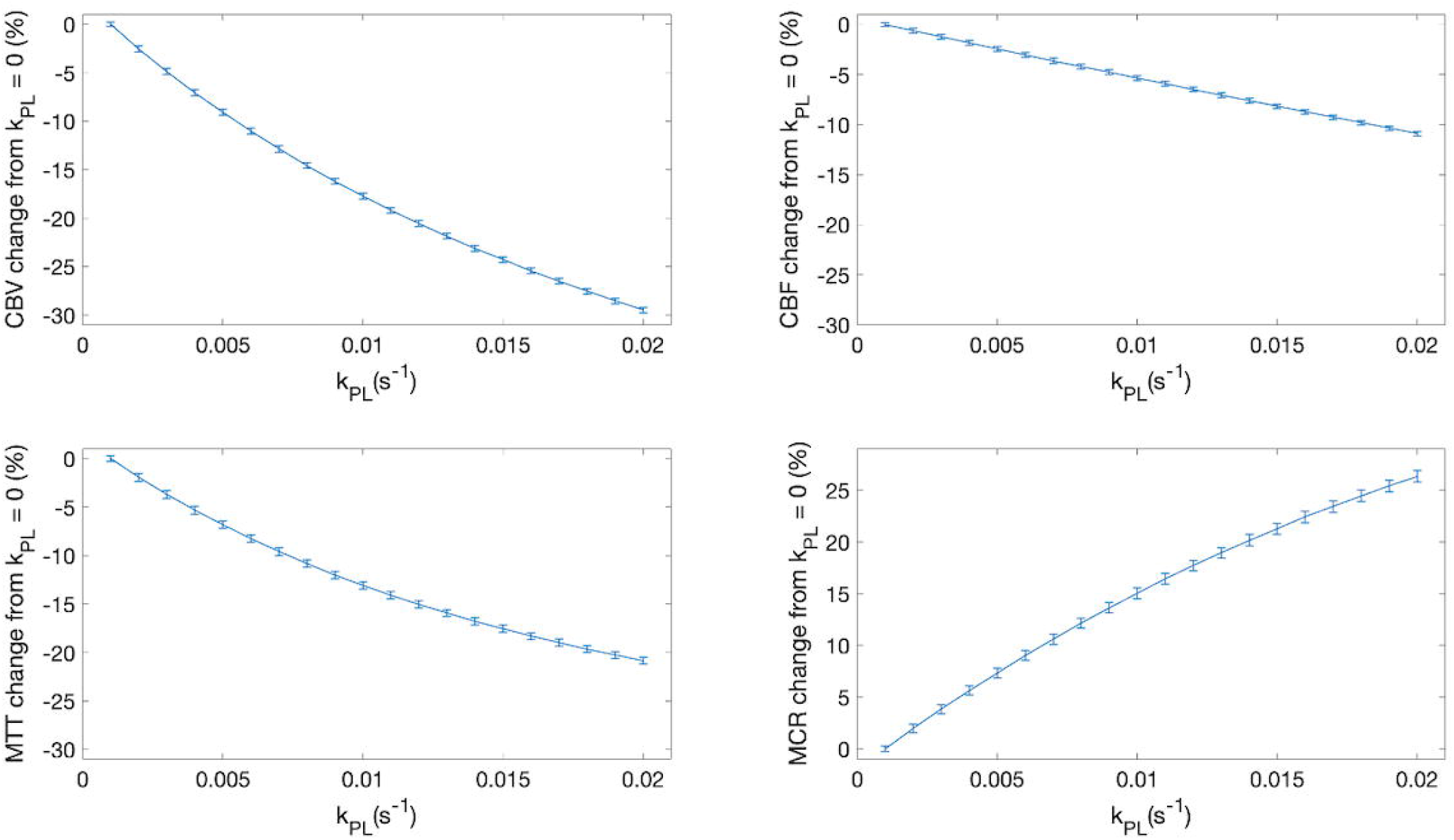
Simulation results demonstrating sensitivity of the model to k_PL_. T_1_pyruvate, B_1+_ deviation, flip angle, and B_1_ error are assumed to be 35s, 1, 10 degrees, and 1, respectively.

**Figure 5.**
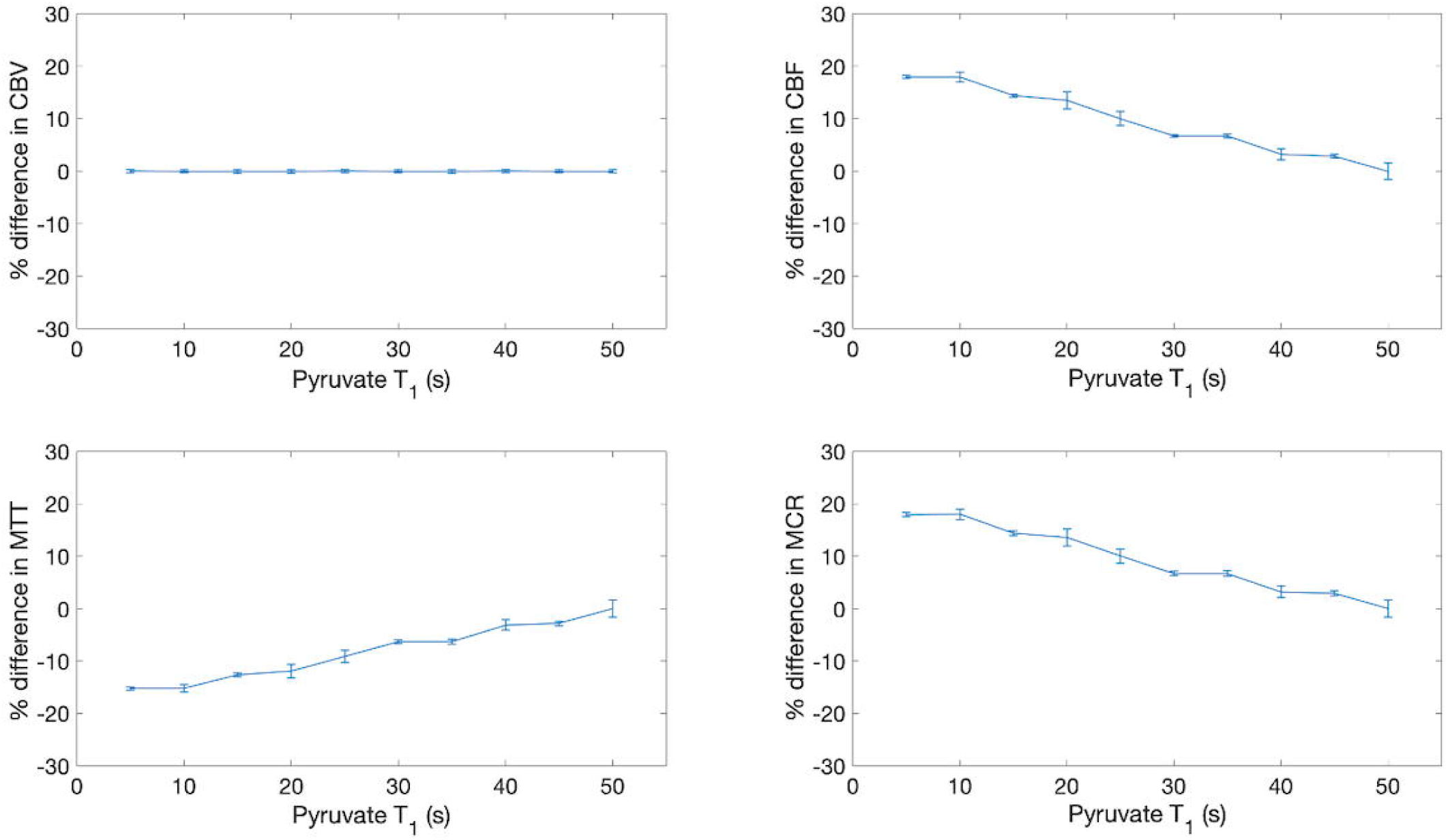
Simulation results demonstrating model variation with pyruvate T1 across all parameters. kPL, flip angle, SNR, and B1 error are assumed to be 0, 10 degrees, 100, and 1, respectively.

Varying the AIF to tissue input function (TIF) size showed very little change in FM when the AIF was much greater (>75x TIF) but high variation in FM as the AIF approached TIF amplitude (FM > 20% at AIF/TIF < 20), see Figure 6.

**Figure 6.**
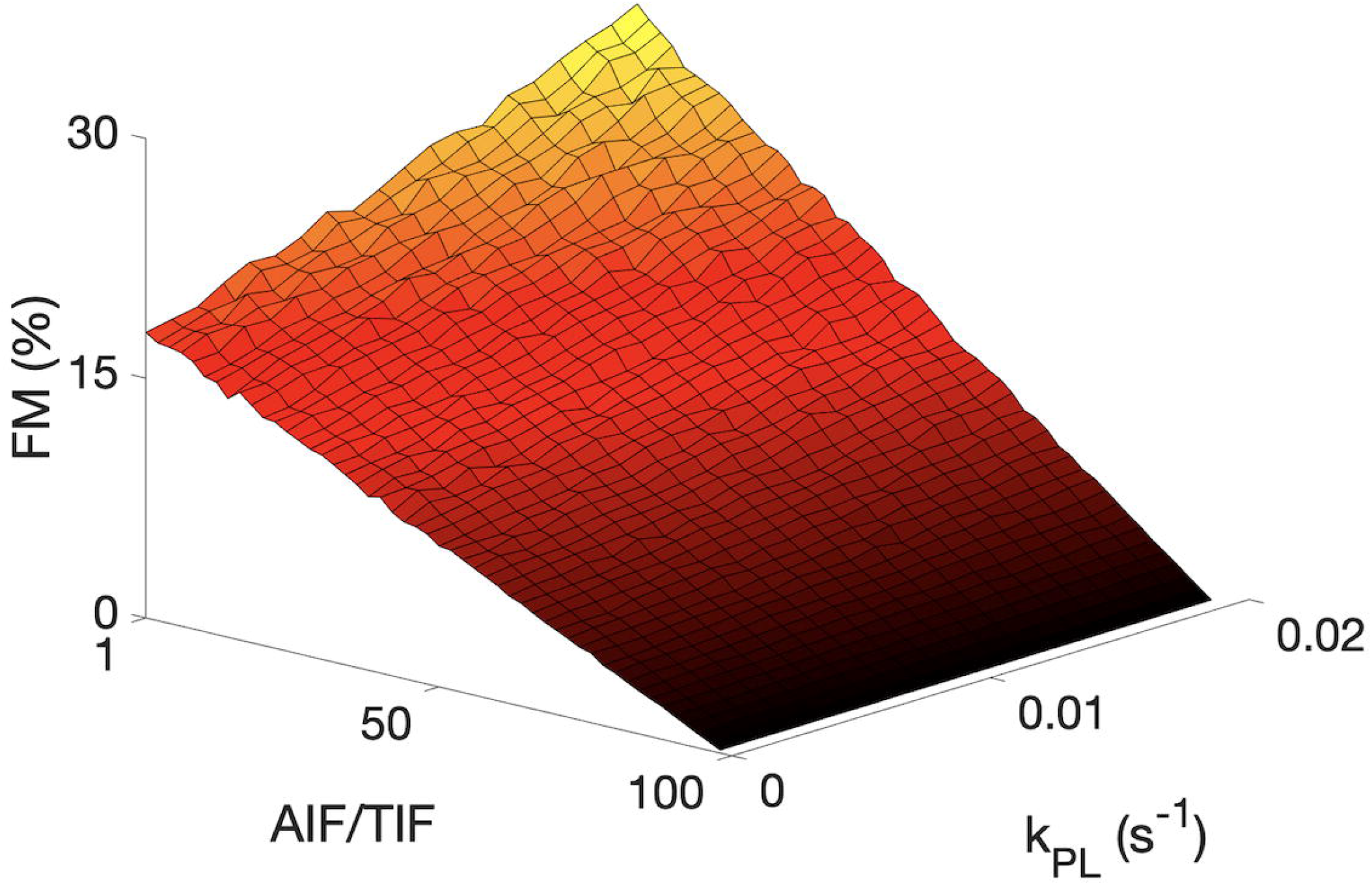
Simulation results showing the variation in FM (%) as both AIF/TIF and kPL are varied. SNR and B1 error are assumed to be 100, and 1, respectively. Generally, an increase in metabolism is observed as perfusion decreased and kPL increased.

### Image analysis

Pre-clinical imaging was successful across species, with variations in metabolic and perfusion parameters seen in the post-stroke brain.

Example post-stroke rodent imaging can be seen in Figure 7 (A to D). There was a significant difference between the ipsilateral and contralateral brain in the rodent model of stroke for CBF_pyruvate_ (140 ± 2 vs 89 ± 6 mL/100g/min, p < 0.01, respectively), CBF_DSC_ (106 ± 8 vs 86 ± 5 mL/100g/min, p = 0.03, respectively), Lactate:Pyruvate (2.7 ± 0.2, vs 2.1 ± 0.2, p = 0.05, respectively) and FM (26 ± 5 vs 4 ± 2 %, p = 0.04, respectively). Detailed results are provided in Table 1.

**Table 1.** Results from ROI analysis from the rodent model of stroke.

| Feature | Ipsilateral | Contralateral | P-value |
| --- | --- | --- | --- |
| TTP<sub>Pyruvate</sub> (s) | 4.5 ± 0.6 | 3.9 ± 0.2 | >0.05 |
| TTP<sub>DSC</sub> (s) | 6 ± 0.5 | 7 ± 0.5 | >0.05 |
| CBV<sub>Pyruvate</sub> (mL/100g) | 1.5 ± 0.4 | 0.9 ± 0.01 | >0.05 |
| CBV<sub>DSC</sub> (mL/100g) | 4.8 ± 0.5 | 4.1 ± 0.9 | >0.05 |
| CBF<sub>Pyruvate</sub> (ml/100g/min) | 140 ± 2 | 89 ± 6 | <0.01* |
| CBF<sub>DSC</sub> (ml/100g/min) | 106 ± 8 | 86 ± 5 | 0.03* |
| MTT<sub>Pyruvate</sub> (s) | 6 ± 1 | 6 ± 1 | >0.05 |
| MTT<sub>DSC</sub> (s) | 1.4 ± 0.3 | 0.97 ± 0.04 | >0.05 |
| MCR (s<sup>-1</sup>) | 0.17 ± 0.04 | 0.16 ± 0.01 | >0.05 |
| Lactate:Pyruvate | 2.7 ± 0.2 | 2.1 ± 0.2 | 0.04* |
| FM (%) | 26 ± 5 | 4 ± 2 | <0.01* |

**Figure 7.**
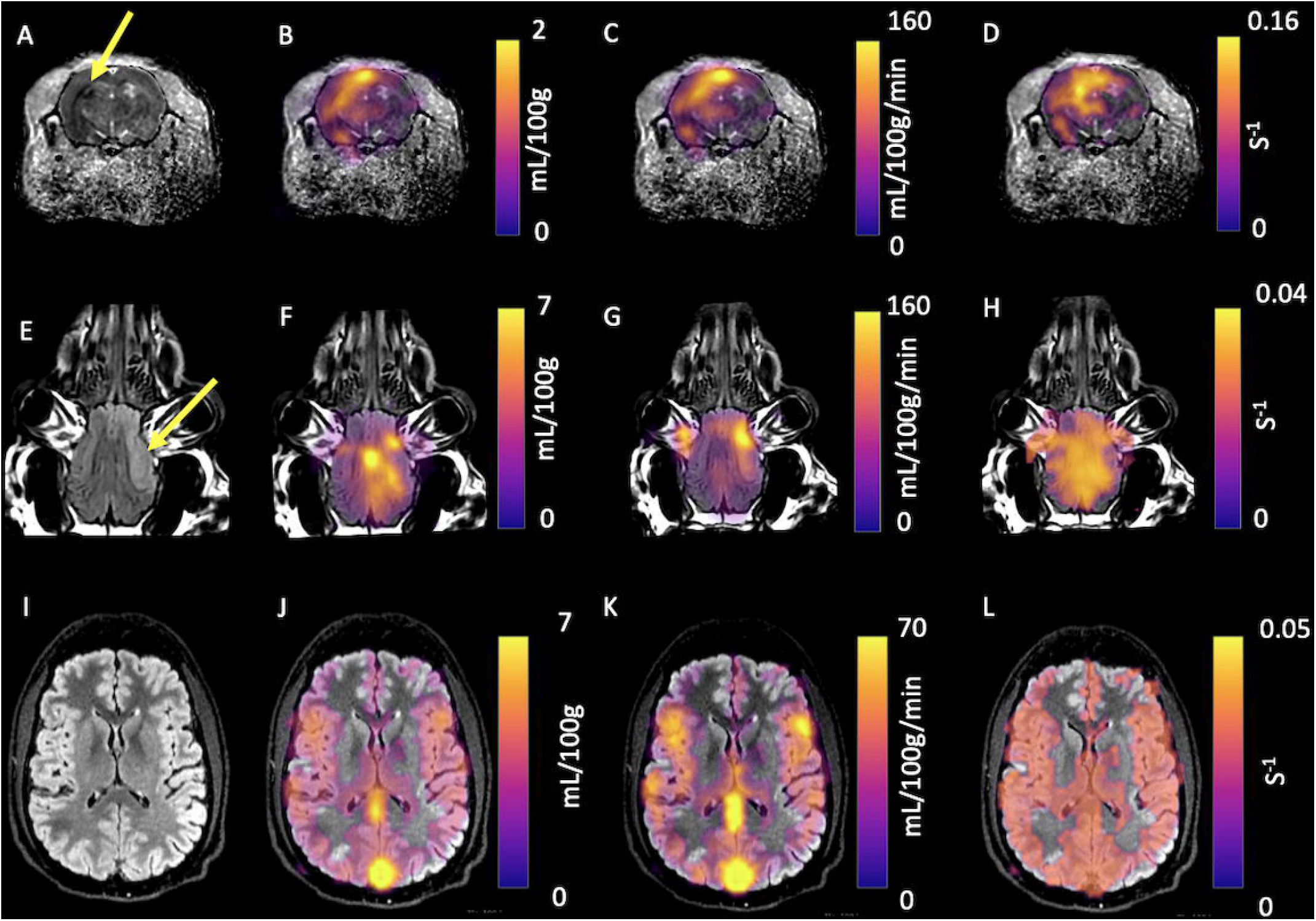
Parametric mapping demonstrated in a rodent and procine models of stroke. The lesion (denoted by the yellow arrow on Apparent Diffusion Coefficient (A) and T_2_ FLAIR weighted imaging (E), with CBV (B and F, respectively), CBF (C and G, respectively), and MCR (D and H, respectively) Also, parametric mapping of the healthy human brain is shown: T_2_ FLAIR (I), CBV (J), CBF (K), and MCR (L).

Example porcine imaging is shown in Figure 7 (E to H). There was a significant difference between ipsilateral and contralateral CBF_pyruvate_ (139 ± 12 vs 95 ± 5 mL/100g/min, p = 0.003, respectively), CBF_DSC_ (116 ± 24 vs 63 ± 40, p = 0.03, respectively), MTT_pyruvate_ (31 ± 8 vs 60 ± 2, p=0.04, respectively), MCR (0.034 ± 0.007 vs 0.017 ± 0.02 s^-1^, p = 0.03, respectively) and Lactate:Pyruvate (1.09 ± 0.07 vs 0.70 ± 0.06, p < 0.01), respectively). Detailed results are provided in Table 2.

**Table 2.**
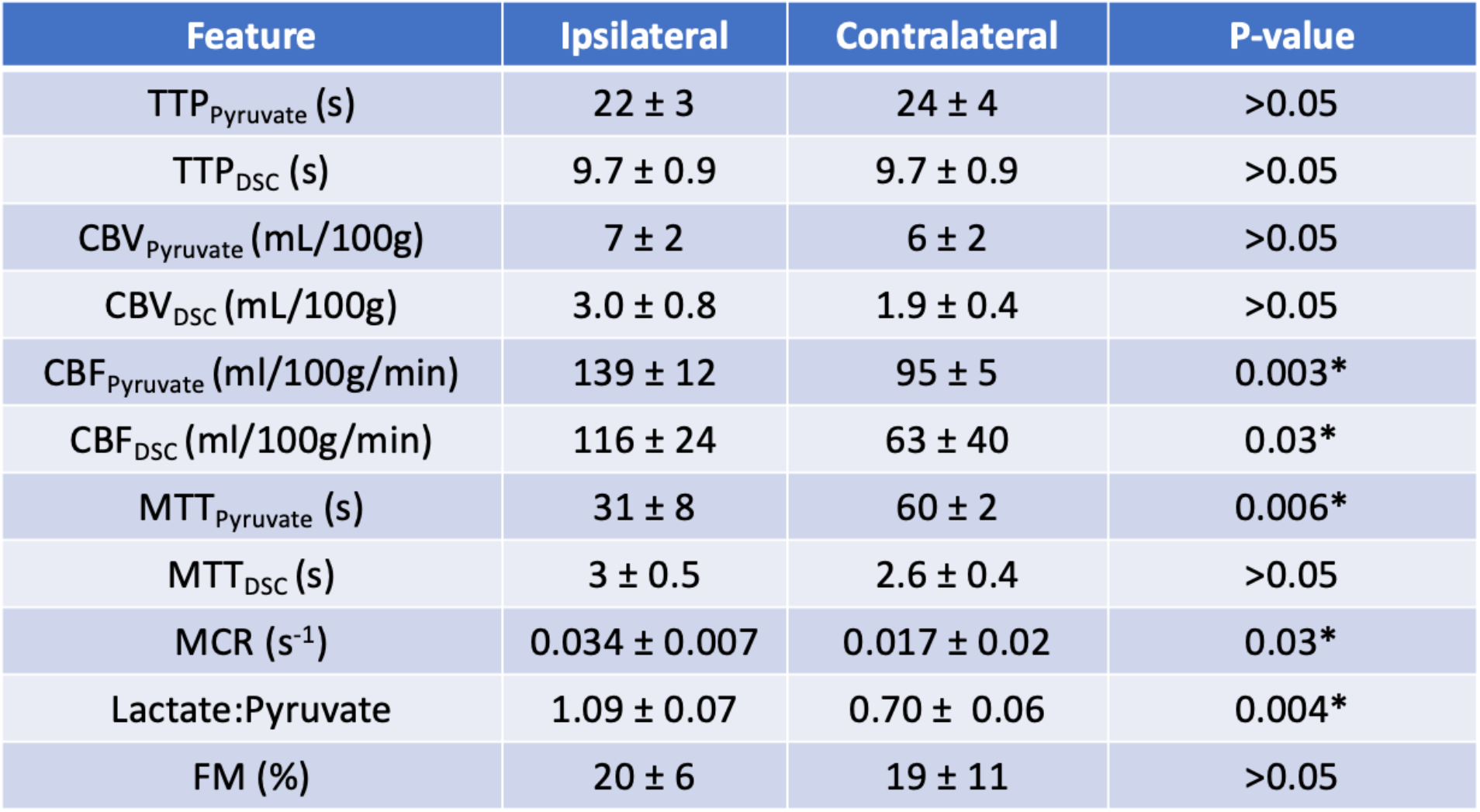
Results from ROI analysis from the porcine model of stroke.

The human imaging was completed successfully, with example images shown in figure 6 (I to L). A significant positive correlation between pyruvate and ASL rCBF was found in the healthy human brain (r = 0.81, p = 0.02), with results shown in figure 8.

**Figure 8.**
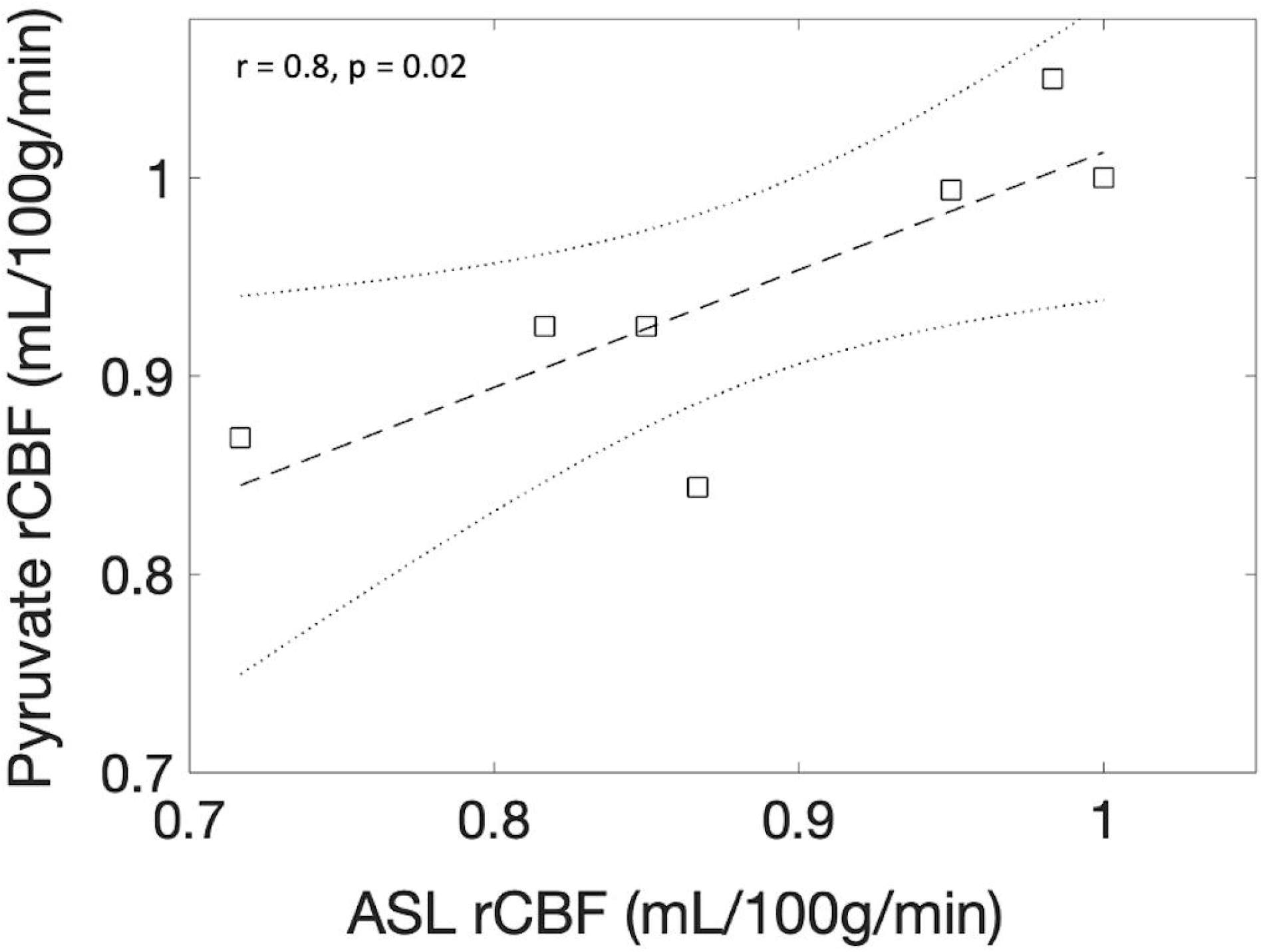
Correlation results from rCBF analysis from the healthy human brain demonstrating a strong correlation between ASL rCBF and pyruvate rCBF.

## Discussion

The main finding in this study is the ability to achieve a quantitative analysis of hyperpolarized ^13^C metabolic imaging data using the metabolic clearance rate formalism utilized in PET^27^. This method is readily implemented, allows for future comparison across sites and between species as demonstrated in both a porcine and rodent model of stroke, and thus supports the progress towards human application in the clinical setting. The method is simple, and the simulation shows that is both reliable and reproducible across a number of potential perturbations to the assumed model.

The metabolic clearance rate formalism extends the sensitivity and specificity of hyperpolarized MR by adding additional information and thus supports additional contrasts for multi-parametric mapping. The fact that the supra-physiological pyruvate signal seems to reflect cerebral perfusion also supports the use of the perfusion measures directly in the analysis. Indeed, a further advantage of using the pyruvate signal is that high-resolution imaging can be performed due to the relatively strong signal from the substrate, in comparison to its downstream low-signal metabolites^28^. Further comparison of model-free deconvolution with modelled solutions using gamma-variate functions could be performed^29^, however the use of model-free deconvolution imposes fewer restrictions on the outcome and thus supports the translation of the method.

The introduction of this methodology to the field of hyperpolarized ^13^C MRI provides quantitative imaging that is akin to that already used in the clinic, notably derived from Arterial Spin Labeling or gadolinium-enhanced MRI^30,31^. Indeed, this approach may provide to be a translation steppingstone for adoption into the clinic, with a focus on imaging outputs that are similar to those already produced and reported. Indeed, the quantitation of CBV and CBF by gadolinium-enhanced imaging is already extensively undertaken in stroke and brain tumors^20,32^, with perfusion known to play a key role in understanding tissue recruitment to the ischemic core (in the case of stroke) or tumor aggressiveness^33^. Of note, the absolute difference in calculated CBF and CBV between perfusion-weighted and hyperpolarized imaging may be derived from the difference in bolus size, leading to a difference in stress response and/or bioavailability of the substrate in comparison to gadolinium, and this warrants further investigation.

The main limitation of this method is the need for a full sampling of the time course of the hyperpolarized tracer, limiting the window of opportunities for selective acquisition schemes, such as chemical shift imaging (CSI), commonly acquiring only one single time point. However, the need to only acquire the injected high SNR substrate simplifies and improves the experimental protocol significantly and combines well with routine clinical perfusion imaging, which is routinely collected in a number of pathologies. Furthermore, most clinical sites are currently opting for dynamic acquisition protocols for hyperpolarized MRI, and so this may not prove to be a large barrier to community adoption of this methodology. Indeed, further validation of this approach to post-processing and quantification of hyperpolarized ^13^C data is needed, especially using multi-site data within and beyond the brain. Indeed, it is noted that this method does not explicitly distinguish between the downstream pathways that are changed due to pathology, for example, elevated [1-13C]Lactate labeling or changes in ^13^C Bicarbonate production but provides potentially complementary information to more targeted measures (such as Lactate:Pyruvate ratio). The variability in measurements of metabolism using different acquisition approaches is yet unknown and is a key hurdle to surpass for the further adoption of this imaging method.

Further investigation and assessment of the added clinical benefit of the MCR and FM formalism, in comparison to *k*_PL_ and ratiometric measures of metabolism, is needed – and this will be a vital component for future clinical radiological studies using hyperpolarized MRI. An additional assessment of the repeatability and reproducibility of the model in both pre-clinical and clinical studies is also needed. Indeed, the actual added clinical value of the technique needs to be further studied, and there is a promise for the method to detect early response to therapy in Breast Cancer^34^, amongst other diseases. Finally, it is of note that this study did not determine the cause of the increase of metabolism in the rodent brain post-stroke; however, it could be due to activated microglia or astrocytes^35^.

## Methods

### Simulations

All simulations were performed in Matlab (The Mathworks, MI, 2019a). Initially, the hyperpolarized pyruvate time course was simulated using a step function arterial input function (AIF) with ongoing radiofrequency and metabolic (*k*_PL_) and T_1_ mediated decay for 200 seconds.

Simulated data were processed using a Matlab implementation of model-free deconvolution^36^, assuming regularization of 0.1. All simulations were run 1000 times per perturbation (perturbations described below).

To understand model stability, model input variables were perturbed one at a time, and the resulting Cerebral Blood Flow (mL/100g/min), Cerebral Blood Volume (mL/100g), Mean Transit Time (s) and metabolic clearance rate (MCR - s^-1^) error (compared to non-perturbed data) was calculated as per equation 4.

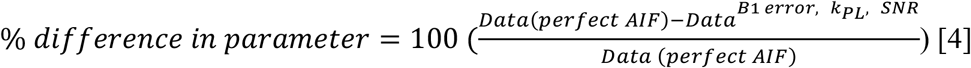

Data (perfect AIF) assumes flip angle = 10 degrees, T_1_ = 35s, Signal to noise ratio (SNR) = 100, Transmit B_1_ error = 1, and *k*_PL_ = 0.

### AIF perturbations

Perturbations included: random noise was added to each time course to give SNR (as defined as the peak signal of the pyruvate curve divided by the standard deviation of the noise at the end of the experiment) varying from 1 to 100. *k*_PL_ was varied from 0 to 0.04 (s^-1^) in steps of 0.001 (s^-1^) (here the tissue concentration of pyruvate was assumed to be 100 times lower than the AIF (extracellular) pyruvate concentration). Transmit B_1_ deviation was varied from 0.1 to 2x nominal flip angle (two simulations: 10 and 30 degrees) in steps of 0.1.

Simulations compared calculated CBV, CBF, and MTT for each *k*_PL_ step to *k*_PL_ = 0 data as per equation 4.

A final simulation was run to assess the sensitivity of FM calculations using a non-metabolized AIF that was scaled in size from 100x Tissue Input Function (TIF), pyruvate in the intracellular space, amplitude to the AIF amplitude. Simultaneously, *k*_PL_ was varied as above for the TIF. The AIF and TIF were used to calculate the FM.

### Pre-clinical experiments

#### Animal experiments

All rodent experiments were conducted in accordance with the XXX regulations. 3 male Wistar rats were anaesthetized with isoflurane, and heads were fixed in a stereotaxic frame. A burrhole was drilled (coordinates relative to bregma, AP +0.1, ML −0.3, SI −0.4 mm), and a fine glass capillary was inserted into the striatum. 1µl of Endothelin-1 was injected over 2 minutes, with the capillary withdrawn and the scalp sutured. Rodents were allowed to recover with food and water ad libitum for 3 days prior to imaging.

Pigs (n = 3, 36-44 kg) were subjected to stroke after approval from the Danish animal inspectorate. The pigs were anaesthetized with propofol and fentanyl. A burrhole was drilled (1.5 cm from the midline just behind the orbit), and 30 µg ET-1 in 200 µl saline was injected over 10 minutes through a 30-gauge needle^37^. The animals were then imaged after 3-4 hours.

#### Hyperpolarization protocol

[1-^13^C] EthylPyruvate (Cambridge Isotopes) was polarized. Briefly, a batch of [1-^13^C] Ethylpyruvate was mixed with AH111501 (Syncom, NL) to a concentration of 15mmolL^-1^. 15uL of Dotarem dissolved in ethanol (2:100 % volume to volume) was added to the batch. 90uL of Ethyl pyruvate was polarized for 45 minutes in a prototype polarizer (Alpha System, Oxford Instruments) prior to dissolution with 4.5mL of sodium hydroxide as previously described ^38^. 1mL of hyperpolarized solution was injected over approximately 4 seconds with a 300uL saline chase.

#### Rodent imaging

Rodent imaging was performed using a 7T MRI system (Agilent magnet). Rats were anesthetized on day 3 post-surgery with 2.5% isoflurane and 60:40% O_2_:N_2_O)^26^, a tail vein cannula inserted, and placed in a cradle with a 2 channel ^13^C receive array (Rapid Biomedical, Rimpar, Germany). Animals were maintained at 2% isoflurane in the magnet with heated air to maintain core temperature. Proton localizers and a 3D RF and gradient spoiled gradient echo (Field of view (FOV) = 60mm^3^, acquisition matrix = 256×256×64, reconstruction matrix = 512×512×64, repetition time (TR) = 4 ms, Echo time (TE) = 2 ms, flip angle (FA) = 10 degrees, averages = 2) imaging was performed using the ^1^H/^13^C body coil for transmit and receive. ^13^C imaging was performed as previously described^38^. Briefly, a multi-echo spiral with interleaved free induction decay (FOV = 40mm^2^, TR (per echo time) = 500ms, FA = 15, acquisition matrix = 40×40, reconstruction matrix = 128, slice thickness = 20mm, Temporal resolution = 4s, total number of time steps = 30) was acquired with iterative decomposition using least squares estimation (IDEAL) from the time of injection as previously described^38^. Images were reconstructed using explicit calculation of the Fourier matrix and a non-uniform fast Fourier transform.

Animals were transferred to a separate cradle for further proton imaging using a 4-channel receive array (Rapid Biomedical, Rimpar, Germany). The cradle featured integrated anaesthetic gas delivery, rectal thermometry, MR-compatible electrical heating, and respiration monitoring, based on designs described elsewhere^39–41^.

The protocol included 3D RF and gradient spoiled gradient echo (as above but with FOV = 40mm^3^, TR = 5.71ms, TE = 2.87ms, FA = 12 degrees), 2D T_2_ weighted imaging (FOV = 40mm^2^, TR = 2000ms, TE = 30ms, echo train length = 2, averages = 4, matrix = 128×128, slice thickness = 0.62mm, reconstruction matrix = 256×256), diffusion weighted imaging (FOV = 40mm^2^, TR = 2000ms, TE = 30ms, echo train length = 2, averages = 4, matrix = 128×128, slice thickness = 0.62mm, reconstruction matrix = 256×256, b values = 0, 1000), and perfusion weighted imaging (FOV = 35mm^2^, matrix = 128×64, slice thickness = 0.62mm, TE = 10ms, TR = 20ms, flip angle = 20 degrees, temporal resolution = 1s, timepoints before injection of contrast. = 3, gadolinium agent = Dotarem, bolus volume = 200uL).

#### Porcine imaging

The porcine data were acquired on a 3T system (MR750, GE Healthcare, WI). A commercial flexible array coil was used for proton imaging (GE Healthcare, WI), while a home-built 14-channel receive coil combined with a commercial transmit (RAPID Biomedical) was used for carbon imaging. After injection of hyperpolarized [1-^13^C] pyruvate (0.875 ml/kg at 5 ml/s of 250 mM pyruvate, 20 ml saline chase) in the femoral vein, spectral-spatial imaging was performed (6°/37° flip angles on pyruvate/metabolites). A stack-of-spirals readout was employed for four slices with 10×10×15 mm^3^ resolution. The temporal resolution was 960 ms for pyruvate. T_1_- and T_2_-FLAIR weighted images were acquired for anatomical reference & a dynamic susceptibility contrast using 2D gradient echo planar imaging exam performed (TR = 800 ms, TE = 25 ms, flip angle = 30 degrees, FOV = 240 mm^2^, slice thickness = 4 mm, slice gap = 5 mm, acquisition matrix = 128×128)

#### Human imaging

After informed consent and approval from the Committee on Health Research Ethics for Central Denmark and the Danish Medicines Agency, a volunteer underwent MRI with hyperpolarized [1-^13^C] pyruvate (male, 38 years). The examination was performed on a 3T scanner (MR750, GE Healthcare, WI) using a dual-tuned ^1^H/^13^C birdcage transceiver coil. Hyperpolarized [1-^13^C] pyruvate was produced according to good manufacturing practice and injected in the antecubital vein (0.43 ml/s at 5 ml/s of 250 mM pyruvate, 20 ml saline flush). A spectral-spatial excitation (flip angles = 12°/70°, TR = 500 ms, TE = 10 ms) was combined with a variable-resolution spiral readout (8.75×8.75 mm^2^ for pyruvate, 17.5×17.5 mm^2^ for metabolites). Six slices of 20 mm were acquired with a temporal resolution of 2 s. In addition to ^13^C-imaging, T_1_-weighted images were acquired for anatomy, and a pseudo-continuous arterial spin labeling sequence was employed for assessment of cerebral blood flow (3D spiral, 3.6×3.6×3.6mm^3^, 2025 ms post label delay, TR = 4.5 s, TE =9.8 ms).

#### Image post-processing and analysis

Perfusion weighted and hyperpolarized imaging were processed with model-free deconvolution^42^ using block-circular deconvolution with in-house reconstruction pipelines in Matlab (2019a, The Mathworks, USA). Voxels below a signal (defined as the summed signal from the ^13^C EthylPyruvate or pyruvate imaging) to noise (defined as a region outside of the brain and body) ratio of 5 were excluded. A region of interest (ROI) was drawn in the supplying vessels of the rat, pig and clinical exam by a physicist with 9 years of experience in neuroimaging.

3D gradient echo images were used for image co-registration in the rodent case. The signal from the [1-^13^C] Ethyl pyruvate and [1-^13^C] pyruvate were used for rodent and porcine/human processing, respectively. The final images consisted of CBV, CBF, MTT, Time to peak (TTP), and MCR. Perfusion-weighted imaging processing was performed in the same manner to hyperpolarized imaging, with signal defined from the pre-contrast injection images, with CBF, CBV, MTT, and TTP maps calculated. Images were overlaid on T_2_ weighted imaging for all studies. ROIs were drawn in the ipsilateral and contralateral hemispheres for both the porcine and rodent imaging, using the perfusion-weighted imaging and T_2_ for guidance, and used for all subsequent statistical analysis.

For the clinical exam, relative CBF results, using an ROI placed in deep grey matter to normalize data, were calculated from ASL and [1-13C] Pyruvate imaging for the healthy human brain. ROIs were placed in various regions of deep grey matter and the spinal column.

#### Statistical analysis

Statistical analysis was performed in R (4.1.0, The R project). The difference in the metabolic and perfusion weighed imaging parameters between ipsilateral and contralateral regions in the rodent and porcine brain was calculated using the Wilcoxon test. The correlation between ASL and pyruvate rCBF was calculated using a linear model. A p-value below 0.05 was considered significant.

## Acknowledgements

James Grist is funded by an EU Horizon 2020 Grant “Alternatives to Gd”, Grant/Award Number: 858149, Damian Tyler would like to acknowledge the British Heart Foundation (Grant/Award Number: FS/19/18/34252), Christoffer Laustsen would like to acknowledge the Lundbeck foundation.

## Author contributions

Study design: JTG, CL, NB

Data acquisition: JTG, YC, NB, AMS, RH, VB, SS

Data processing: JTG, NB

Statistical Analysis: JTG

Manuscript preparation and review: JTG, NB, ESSH, AMS, RH, VB, JJJJM, SS, YC, AB, DJT, CL.

## Data availability

All data are available upon request to the corresponding author.

## Additional information

There are no competing interests to declare.

