## Supporting Figure 1 for "Metabolic clearance rate modeling: A translational approach to quantifying cerebral metabolism using hyperpolarized [1-13C]pyruvate"

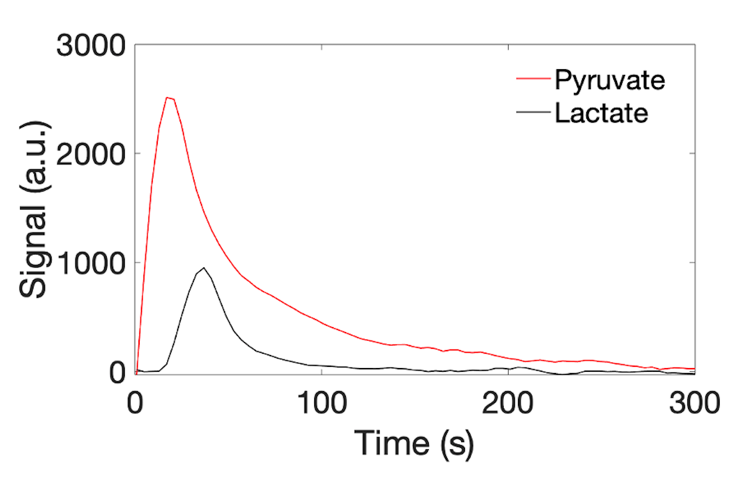

Example simulated metabolic curves for intracellular pyruvate (red) and lactate (black) . Here T1Pyruvate, T1Lactate, TR, kPL, and SNR are assumed to be 35s, 30s, 4s, 0.012s-1, and 100, respectively.
